# Cryo-EM structure of the human CST•Polα/Primase complex in a recruitment state

**DOI:** 10.1101/2021.12.17.473235

**Authors:** Sarah W. Cai, John C. Zinder, Vladimir Svetlov, Martin W. Bush, Evgeny Nudler, Thomas Walz, Titia de Lange

## Abstract

The CST•Polα/Primase complex is essential for telomere overhang maintenance and additionally functions to counteract resection at double-strand breaks. We report a 4.6-Å resolution cryo-EM structure of CST•Polα/Primase, captured prior to catalysis in a recruitment state, which provides insights into the architecture and stoichiometry of the fill-in machinery. Our model informs on human disease mutations that cause Coats plus syndrome.

## Main

Human telomeric DNA terminates in a 3’ overhang of the G-rich strand, which is required for t-loop formation and telomere protection^1^. The mature 3’ overhang must be regenerated during each cell cycle in a controlled manner, and dysfunction of the factors involved can lead to rapid telomere shortening and related human disease^2–4^. Following replication, nucleolytic resection of the 5’ strand can result in excessively long overhangs. This is counteracted through fill-in DNA synthesis by the Ctc1-Stn1-Ten1 (CST) complex and DNA Polymerase α (Polα)/Primase, which are recruited to telomeres by the shelterin complex^2,5–9^ (Fig. 1a-b). In addition to its telomeric function, CST•Polα/Primase performs an analogous fill-in reaction at resected double-strand breaks (DSBs)^10,11^. Thus far, molecular details of how CST interacts with Polα/Primase are largely unknown^12,13^.

**Figure 1:**
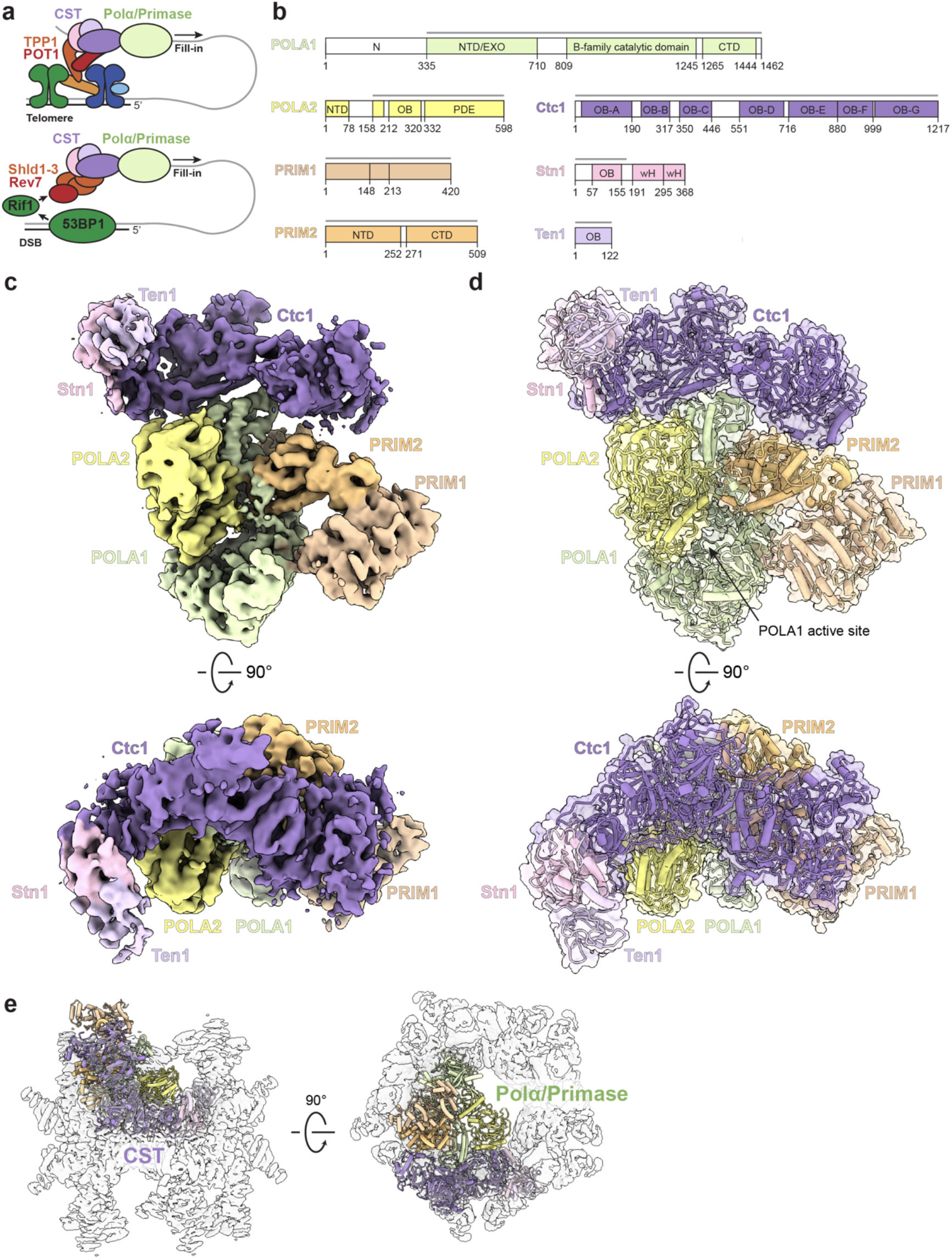
Architecture of the human CST•Polα/Primase complex. **a**, Cartoon schematic of fill-in DNA synthesis orchestrated by CST and Polα/Primase at telomeres (top) and a double-strand break (DSB) (bottom). **b**, Domain schematics for proteins used in this study. Colored bars indicate regions modeled in **d**. **c**, Gaussian-filtered orthogonal views of the cryo-EM map of CST•PP^ΔN^, segmented and colored by subunit. **d**, Atomic model of CST•PP^ΔN^ shown in cartoon and surface representations. **e**, CST•PP^ΔN^ structure (colored cartoon representation) docked into the cryo-EM map of the CST decamer (transparent white surface).

We sought to structurally characterize the complex using recombinant human CST and Polα/Primase purified from insect cells. Although we could reconstitute a stable complex of CST•Polα/Primase using size-exclusion chromatography (SEC), the complex appeared to dissociate in negative-stain EM (Extended Data Fig. 1a-b). We added a [GGTTAG]_3_ substrate, as telomeric ssDNA was previously shown to stabilize CST^14^, and used GraFix^15^ to cross-link the complex, resulting in a higher proportion of intact complexes in cryo-EM 2D averages (Extended Data Fig. 1c-d). We could unambiguously identify CST and Polα/Primase in the 3D reconstructions, but the substantial heterogeneity present in the dataset limited the resolution to >17 Å (Extended Data Fig. 2).

The POLA1 subunit of Polα/Primase contains a disordered N-terminal region (POLA1^N^, a.a. 1-335) that is dispensable for catalysis and omitted in most structures of the enzyme^16^. We purified Polα/Primase lacking POLA1^N^ (referred to hereafter as PP^ΔN^) and reconstituted a CST•PP^ΔN^(•ssDNA) complex as we did with full-length Polα/Primase (referred to hereafter as PP^FL^) and cryo-EM data (Extended Data Fig. 3a). The omission of POLA1^N^ resulted in lower CST occupancy, so we introduced additional classification steps to select for particles containing intact CST (Extended Data Fig. 3c). 131,850 particles were used to generate our final map with a global resolution of 4.6 Å, although local resolution estimates revealed lower resolution for the peripheral regions of CST and PP^ΔN^, likely due to flexibility (Fig. 1c, Extended Data Fig. 3d, e).

The crystal structure of *apo* PP^ΔN^ (PDB ID: 5EXR)^16^ and the cryo-EM structure of CST (PDB ID: 6W6W)^14^ could readily be docked into our density. We then substituted the Ctc1 structure with an AlphaFold model^17,18^, which provides information about the Ctc1 N-terminus that was poorly resolved in the CST cryo-EM map^14^. After initial rigid-body docking followed by flexible fitting and refinement, the overall conformations of the two subcomplexes did not show major changes from their structures in isolation. Only the C-terminal domain of POLA1 (POLA1^CTD^) moves slightly to contact Ctc1 (Fig. 1d, 2a). From the overall architecture of the complex, we can draw three conclusions.

**Figure 2:**
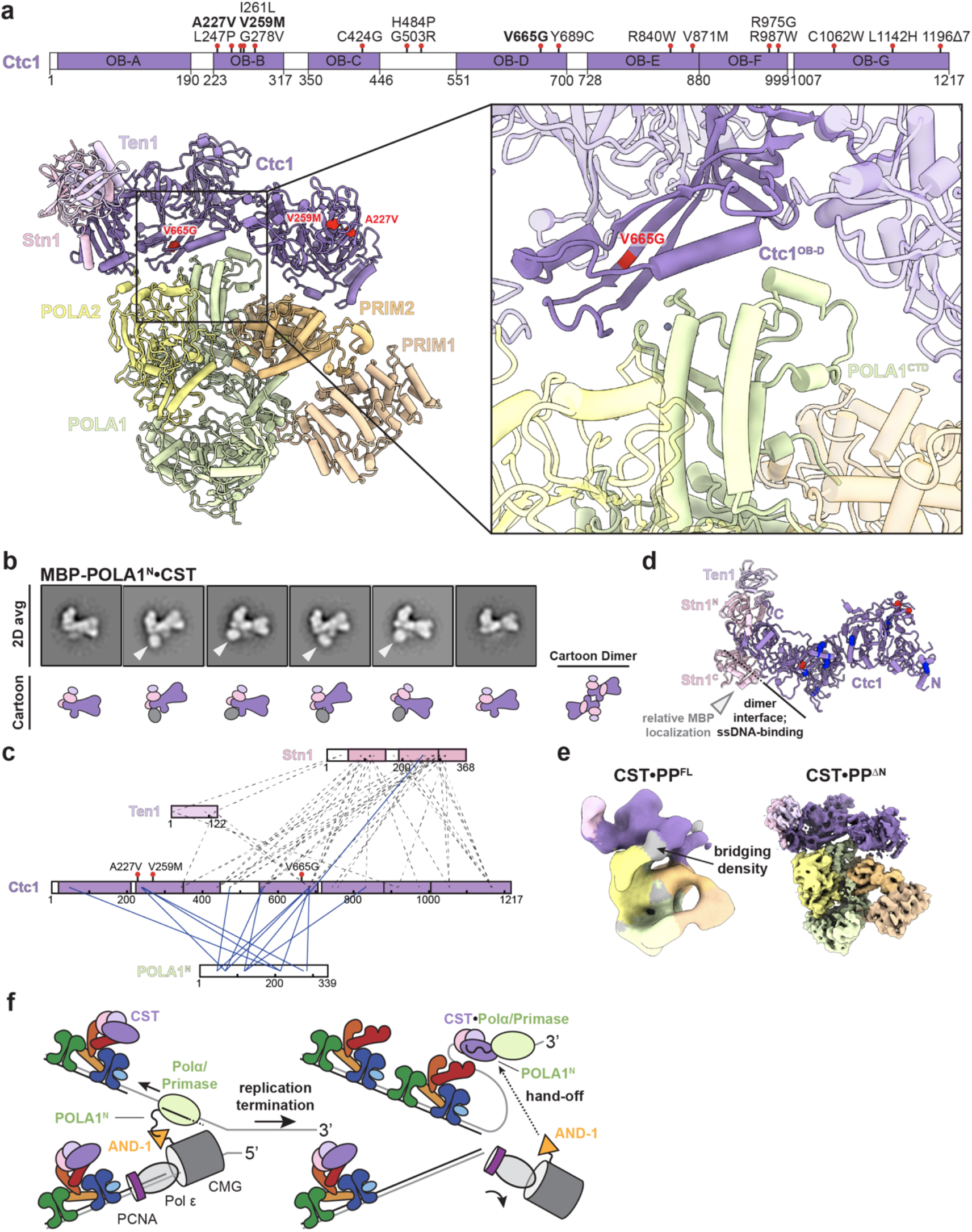
Details of the CST•Polα/Primase interaction and CP mutations. **a**, Summary of Coats plus point mutations in *CTC1* and mapping of Polα/Primase-binding mutations (highlighted in bold face) on the CST•PP^ΔN^ structure. Box: zoomed-in view of the primary Ctc1^OB-D^-POLA1^CTD^ interaction. **b**, Averages and cartoon representations of the six most populated *RELION* 2D classes of the MBP-POLA1^N^•CST complex. The MBP mass label is indicated with a white arrowhead when present. **c**, Cross-links between POLA1^N^ and CST subunits identified by CX-MS. Blue lines: cross-links between CST and POLA1^N^. Dashed lines: inter-subunit cross-links in CST. **d**, Cartoon representation of the CST monomer in the same view as in **b**. Lysine residues cross-linked to POLA1^N^ (**c**) are shown as blue spheres, and the three CP mutations mapped in **a** are shown as red spheres. White arrowhead indicates relative localization of the MBP mass label. **e**, Comparison of the CST•PP^ΔN^ and CST•PP^FL^ (class 3, Extended Data Fig. 2a) cryo-EM density maps, segmented and colored according to the CST•PP^ΔN^ model. Density observed in the CST•PP^FL^ map not accounted for by the model is shown in grey. **f**, Proposed model for the handoff of Polα/Primase to shelterin-bound CST following telomere replication.

First, in the complex Polα/Primase remains in the auto-inhibited state^16^, where the POLA2 subunit blocks the active site of POLA1 (Fig. 1d). This is consistent with our chemical cross-linking analysis, which showed that cross-linking preferentially stabilizes the more compact, occluded state compared to the flexible, extended state of the enzyme (data not shown)^19^. Thus, we conclude that our structure likely captures a recruitment state of the complex that forms prior to active RNA/DNA synthesis by Polα/Primase.

Second, the CST•Polα/Primase interaction is sterically incompatible with the CST decamer, as it would bind in the center of the ring and sterically clash with neighboring CST subunits (Fig. 1e). CST has also been shown to dimerize upon ssDNA binding^14^, but two lines of evidence suggest that active CST•Polα/Primase has a 1:1 stoichiometry. (1) We do not observe CST dimers in our 2D-class averages (Extended Data Figs. 1, 2). (2) We characterized the CST-POLA1^N^ interaction to understand why CST•PP^FL^ did not yield a high-resolution map although PP^FL^ forms a more stable interaction with CST. We reconstituted a native complex of CST and MBP-tagged POLA1^N^ and analyzed it by negative-stain EM and cross-linking mass spectrometry (CX-MS). With MBP as a mass label, we localized the N-terminus of POLA1^N^ to the primary CST dimerization interface (Fig. 2b). CX-MS analysis shows that POLA1^N^ may bind in multiple modes to CST, which could partially explain the heterogeneity observed with CST•PP^FL^ (Fig. 2c-e, Extended Data Fig. 4).

Third, the primary interaction interface observed in our structure occurs between POLA1^CTD^ and CTC1^OB-D^, which is an elongated OB-fold that shares no homology to known OB-folds^14^ (Fig. 2a). This is particularly interesting given the proposal that CST and Polα co-evolved in eukaryotes with telomeres, as it raises the possibility that CST evolved specificity for Polα through this unique OB-fold^20^. Our structural model also places regions of Ctc1 near POLA2 and PRIM2. These two potential interaction sites are not well resolved in our cryo-EM map, but it is possible that these interactions are weak or more transient as CST flexes about the hinge generated by the POLA1^CTD^/CTC1^OB-D^ interaction (Fig. 1c-d, Fig. 2a).

Dysfunctional fill-in, primarily driven by mutations in CST, causes the severe developmental disorder Coats plus syndrome (CP)^3,4^. Three point mutations (A227V, V259M, and V665G) in Ctc1 were previously described to disrupt Polα/Primase association with CST^12^. We mapped these residues onto our structure to investigate the molecular basis of dysfunction caused by these mutations. V665 resides on a β-strand of CTC1^OB-D^, so it is plausible that a glycine substitution would destabilize the β-sheet and OB-fold, disrupting the primary interaction that we observe. The mutations at A227 and V259 reside on CTC1^OB-B^, which does not contact Polα/Primase in the CST•PP^ΔN^ structure (Fig. 2a). However, one major difference we observe between the cryo-EM maps of CST•PP^ΔN^ and CST•PP^FL^ is the presence of connecting density between POLA2 and the Ctc1N-terminus, which we only see when POLA1^N^ is present. Because POLA1^N^ and POLA2^NTD^ (a.a. 1-78, attached by a flexible linker and not visualized in our CST•PP^ΔN^ structure) have been described to cooperatively bind in other settings^21^, we speculate that this bridging density is a combination of these two features. Although our resolution is limited, this connection could potentially explain the CP mutations occurring at the Ctc1 N-terminus (Fig. 2a, e).

In summary, our cryo-EM structures of CST•Polα/Primase reveal the fill-in machinery captured in an occluded state. We propose that this state occurs during recruitment of Polα/Primase to the telomere, prior to the start of RNA/DNA synthesis by the enzyme. POLA1^N^ is responsible for the flexible tethering of Polα/Primase to the replisome via an interaction with AND-1^22,23^, and its extensive interaction with CST suggests a potential spatiotemporal regulation of fill-in after telomere replication, where the replisome hands Polα/Primase off to a shelterin-bound CST for C-strand synthesis (Fig. 2f).

---

**Extended Data Table 1.**
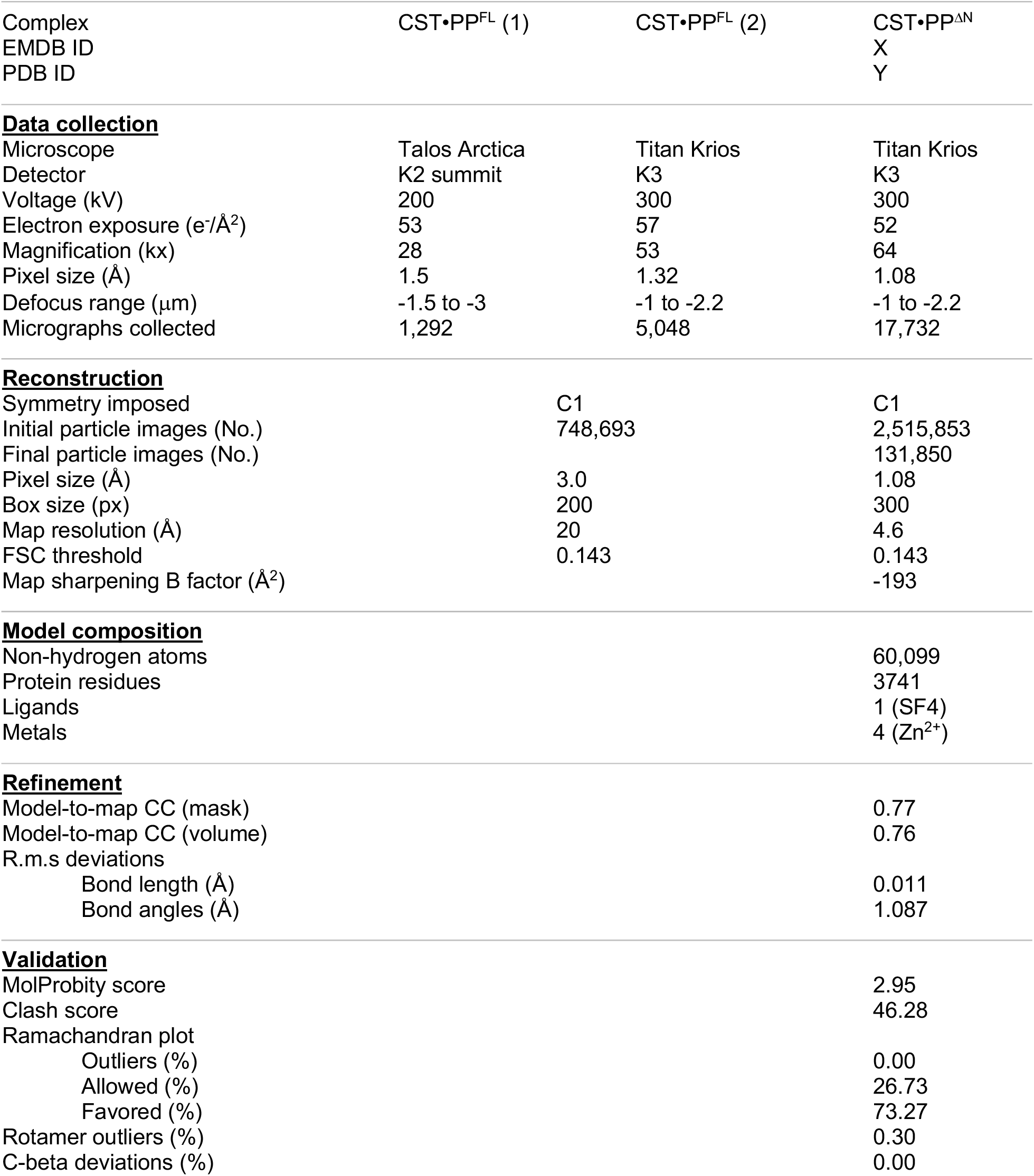
Cryo-EM data collection and validation statistics.

**Extended Data Figure 1:**
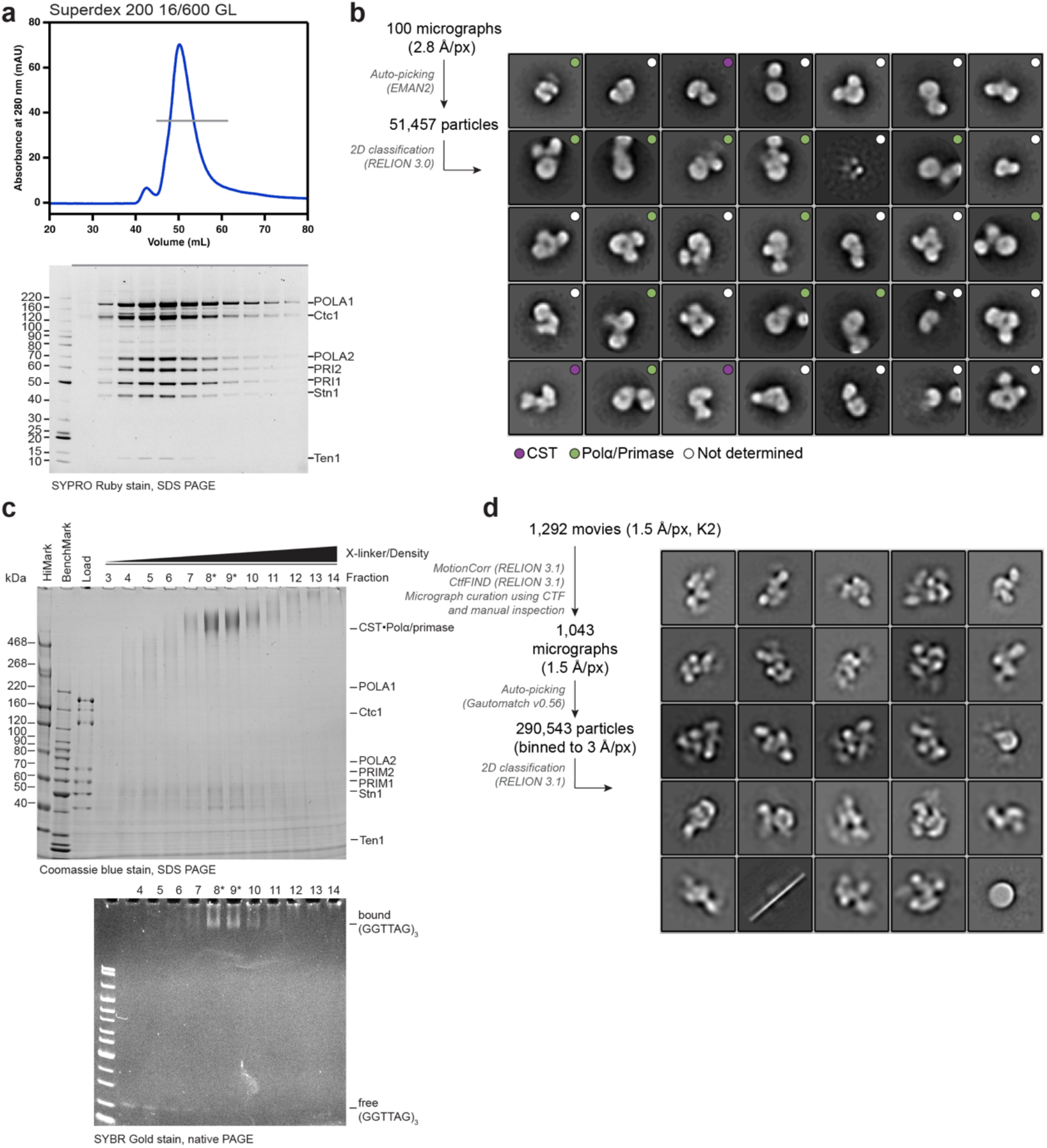
Reconstitutions of CST•PP^FL^. **a**, SEC profile of CST•PP^FL^ (top). The grey line indicates the fractions visualized on the SYPRO Ruby-stained SDS-PAGE gel (bottom). **b**, Negative-stain EM image-processing pipeline and averages of the 35 (of 50) most populated 2D classes of the reconstituted complex characterized in **a**. The size of the 128-px box corresponds to 358 Å. **c**, GraFix preparation of CST•PP^FL^•ssDNA. *Top:* Coomassie blue-stained SDS-PAGE gel (4-12% (w/v) Tris-Glycine) of GraFix fractions, showing formation of cross-linked CST•PP^FL^ species. Fractions marked with an asterisk (*) were pooled for analysis by cryo-EM. *Bottom:* SYBR Gold-stained native PAGE gel (4-20% (w/v) TBE) of GraFix fractions, showing free and bound ssDNA. **d**, Cryo-EM image-processing pipeline and 2D class averages obtained after vitrification and imaging of the GraFixed complex shown in **c**. See Extended Data Table 1 for full data collection and processing details.

**Extended Data Figure 2:**
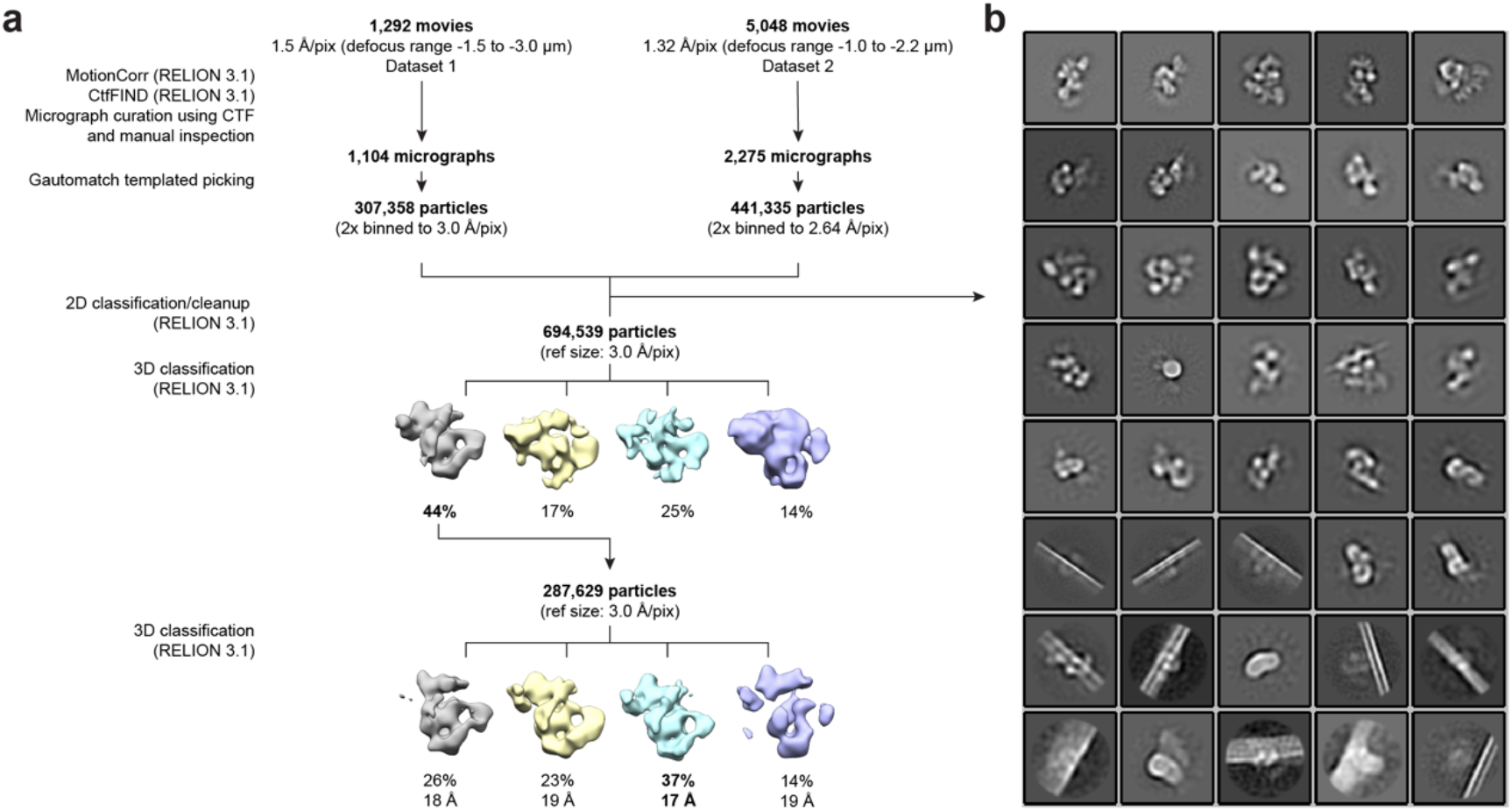
Cryo-EM image-processing pipeline of CST•PP^FL^. **a**, Cryo-EM image-processing pipeline used for the CST•PP^FL^ complex. Two datasets were combined in *RELION 3.1* as separate optics groups, and particles were refined together with a reference generated from dataset 1. See Extended Data Table 1 for full data collection and processing details. **b**, 2D-class averages from a cleanup step prior to 3D classification. Classes clearly representing ice contamination or graphene oxide edges were removed during cleanup.

**Extended Data Figure 3:**
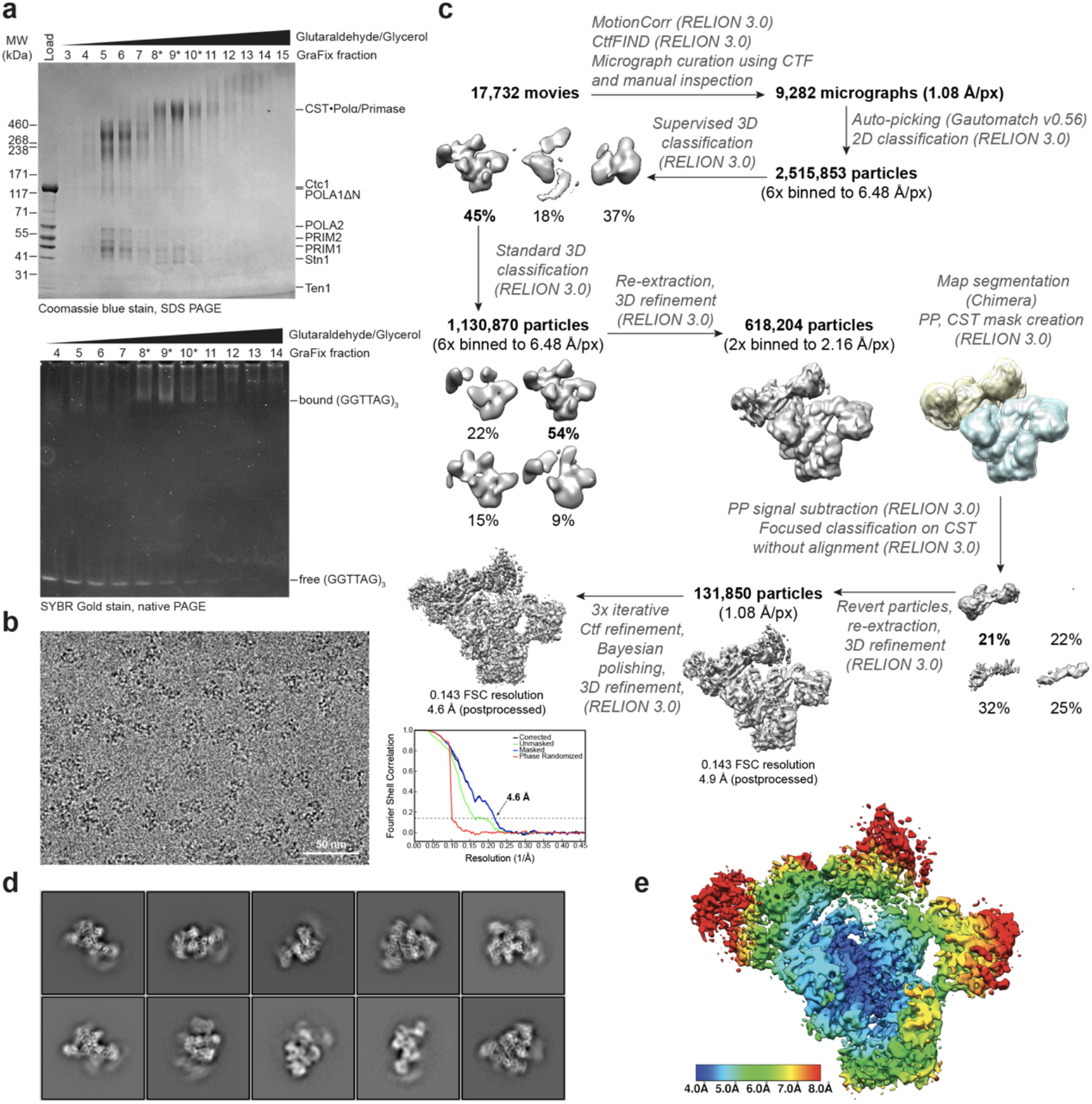
GraFix preparation of CST•PP^ΔN^ complex and analysis by cryo-EM. **a**, GraFix preparation of CST•PP^ΔN^•ssDNA. *Top:* Coomassie blue-stained SDS-PAGE gel (4-12% Tris-Glycine) of GraFix fractions, showing formation of cross-linked CST•PP^ΔN^ species. Fractions marked with an asterisk (*) were pooled for analysis by cryo-EM. *Bottom:* SYBR Gold-stained native PAGE gel (4-20% TBE) of GraFix fractions, showing free and bound ssDNA. **b**, Selected motion-corrected cryo-EM micrograph of CST•PP^ΔN^. **c**, Cryo-EM image-processing pipeline used for the CST•PP^ΔN^ complex, including supervised 3D classification and focused 3D classification steps used to select particles with intact CST. Gold-standard FSC curves for the final map of CST•PP^ΔN^ indicate a global resolution of 4.6 Å (FSC = 0.143). See Extended Data Table 1 for full data collection and processing details. **d**, 2D-class averages obtained with the particle stack (131,850 particles) used to generate the final CST•PP^ΔN^ map show blurring at the peripheral regions, indicating flexibility. **e**, Local resolution estimates of the CST•PP^ΔN^ map.

**Extended Data Figure 4:**
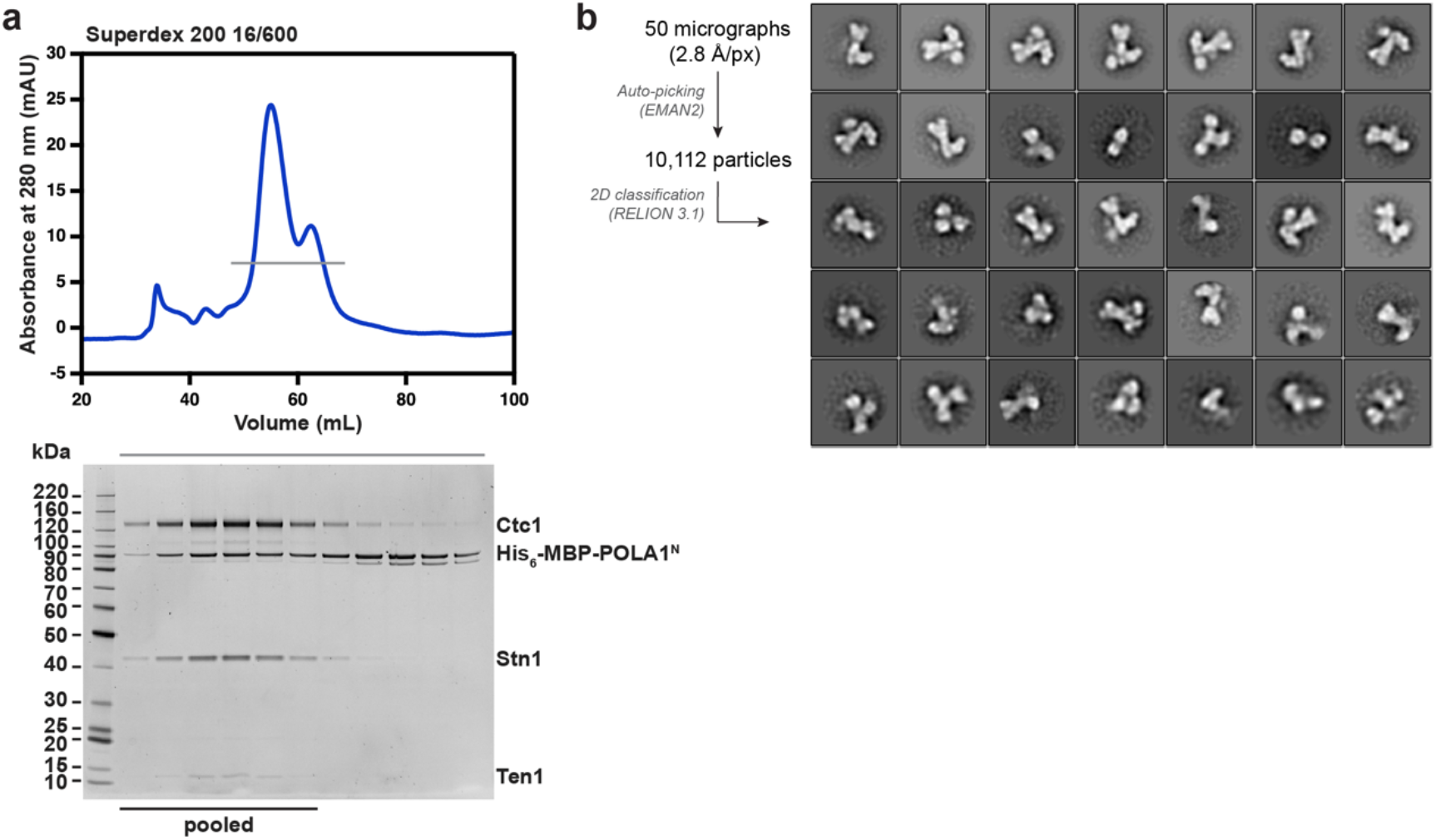
Reconstitution of the MBP-POLA1^N^•CST complex. **a**, SEC profile of reconstituted His_6_-MBP-POLA1^N^•CST (top). The grey line indicates the fractions visualized on the SYPRO Ruby-stained SDS-PAGE gel (bottom). The black bar indicates the fractions pooled for negative-stain EM and CX-MS analysis (see Fig. 2). **b**, Negative-stain EM image-processing pipeline and averages of the 35 (of 50) most populated classes of the reconstituted complex characterized in **a**. The size of the 128-px box corresponds to 358 Å^2^.

## Acknowledgements

We are grateful to H. Suzuki and Y. Zhang for guidance with data processing. We thank M. Ebrahim, J. Sotiris and H. Ng at the Evelyn Gruss Lipper Cryo-EM Resource Center of The Rockefeller University, for assistance with cryo-EM data collection. SWC is supported by the National Science Foundation Graduate Research Fellowship under Grant No. 1946429. JCZ was previously supported by an NIH cancer biology training grant (T32 CA009673) and is currently the Lorraine Egan Fellow of the Damon Runyon Cancer Research foundation (DRG 2337-19). This work is supported by grants to TDL from the NIH (5 R35 CA210036) and the Breast Cancer Research Foundation (BCRF-19-036) and by funding from the Howard Hughes Medical Institute to EN and VS.

## Author contribution

This study was conceived by TdL and TW. CX-MS data was obtained by EN and VS. All other experiments were executed by SWC with help from JCZ. MWB helped with negative-stain EM experiments. SWC wrote the manuscript with input from all authors.

